# Novel action of proline-rich antimicrobial peptides Api88, Api137, Onc72 and Onc112 against *Pseudomonas aeruginosa* and *Escherichia coli* in ion-rich environments

**DOI:** 10.1101/2025.05.09.653031

**Authors:** Namfon Pantarat, Daniel Knappe, Eric C Reynolds, Ralf Hoffmann, Neil M. O’Brien-Simpson

**Author notes:** Correspondence: Neil M. O’Brien-Simpson; Ralf Hoffmann.

## Abstract

The rise in antibiotic resistance has meant that there is a need for new strategies and one avenue is the use of proline-rich antimicrobial peptides (PrAMPs). Here we investigate how different metal ion environments (Na^+^, Mg^2+^, Ca^2+^) affect antimicrobial activity of PrAMPs derived from apidaecin 1b (Api88, Api137) and *Oncopeltus* antibacterial peptide 4 (Onc72, Onc112) against *Pseudomonas aeruginosa* and *Escherichia coli*. Initial antimicrobial testing in an ion-rich media (ion levels similar to mammalian body fluids) found that the PrAMPs were effective against *E. coli* but not *P. aeruginosa*. Both Api88 and Api137 were bactericidal, while Onc72 and Onc112 were bacteriostatic against *E. coli*. In a lower ion-media the activity of the PrAMPs significantly improved against both bacteria and Onc72 and Onc112 altered the mode of action to bactericidal. In low Na^+^, Ca^2+^ and Mg^2+^ ion conditions all of the peptides were able to penetrate the outer membrane of *P. aeruginosa*, however at higher ion concentrations none of the peptides were able to penetrate the outer membrane. PrAMPs were found to cause *E. coli* cells to swell and have a hyperpolarised membrane indicating a new mechanism of action for PrAMPs. Our data indicates that bacteria reduce susceptibility to AMPs by stabilising their LPS layer with metal ions and that the PrAMPs have secondary modes of action affecting the functionality of the bacterial membrane. Combining an ion chelator with PrAMPs may be a novel solution to combat weak antimicrobial activity in ion-rich environments such as host tissues.

## Introduction

By 2050 it is predicted that more people will die from bacterial infections than cancer and this mortality will have a projected economic impact of U.S $60-100 trillion per year^1^. Currently, multidrug resistant (MDR) bacterial infections cause >700,000 deaths/year and incur an estimated annual treatment cost of >U.S. $20 billion^2^. Antimicrobial resistance is considered as *‘one of our most serious health threats’*^2^ and there is now unanimous agreement that new, potent and selective antimicrobial (AM) therapies are urgently required. Although there has been a trend away for developing modified antibiotics due to the rapid induction of resistance there is a class of materials where resistance development is extremely slow and these antimicrobial peptides (AMPs) and in particular proline rich AMPs (PrAMPs) are being developed for clinical use^3^.

Antimicrobial peptides (AMPs) are expressed in all kingdoms of life as part of innate immunity in response to infection. Their evolutionary success in fighting a wide variety of pathogens is due to their large structural diversity and the use of different mechanisms to target, for example, the outer membrane or intracellular targets of bacteria. While the expression of these peptides results in rather low concentrations in the host, therapeutic inventions require a greater margin of safety, as high concentrations are observed shortly after administration and decrease until the next dose. Therefore, many AMPs that target the membrane by a lytic mechanism typically provide a low margin of safety, because they also attack mammalian membranes at higher concentrations. PrAMPs, which are expressed in insects and mammals but not in humans, represent a promising class of lead structures because they target intracellular bacterial targets and are relatively stable against proteolytic degradation due to the high content of proline residues, which significantly reduce or even prevent enzymatic cleavage at their N-terminal site^4, 5^. Insect-derived PrAMPs appear to be interesting lead structures due their intrinsic properties and their relatively short sequences of 18 to 20 residues, which can be chemically synthesized in high yields. Solid phase peptide synthesis is well established and allows the rapid and inexpensive synthesis of many analogues by substitution of canonical to non-canonical amino acids at various positions, including backbone modifications, to improve proteolytic stability, target binding and specificity, and more generally the pharmacokinetic properties of peptides.

In recent years, we have optimised the structures of apidaecin 1b, an 18-residue PrAMP expressed in honeybees, resulting in Api88 and Api137, and *Oncopeltus* antibacterial peptide 4, a 20-residue PrAMP expressed in milkweed bugs, resulting in oncocins Onc72 and Onc112 (Table 1)^4-8^. These peptides were highly effective in several murine infection models, despite rather rapid renal clearance^7-13^. Recent data indicate that native PrAMPs, including our lead compounds, can penetrate the outer membrane of Gram-negative bacteria to reach the periplasm, where they mostly rely on the SbmA transporter to enter the cytoplasm, although some PrAMPs, including Onc112, use the MdtM transporter to enter cells^14^. While the heat shock protein DnaK was first suggested as a binding partner of PrAMPs, later studies identified the bacterial ribosome as lethal target for both insect- and mammalian-derived PrAMPs^15, 16^. They inhibit protein translation by binding to the polypeptide exit tunnel (PET) and disrupting ribosome assembly. Later studies using high-resolution cryogenic electron microscopy (cryo-EM), revealed that Onc112 blocks and destabilizes the initiation complex, whereas Api137 binds to the PET near the peptidyl transferase center and inhibits protein translation, traps release factors and thereby arrests ribosomes at stop codons^17, 18 19,20^. Recently, we identified a second binding site for Api88 and Api137 in the PET and a third binding pocket deep within domain III of the 23S rRNA for Api88^21^ and structurally confirmed our previous hypothesis that Api137 disrupts the assembly of the large (50S) subunit of bacterial ribosomes^22^.

**Table 1.**
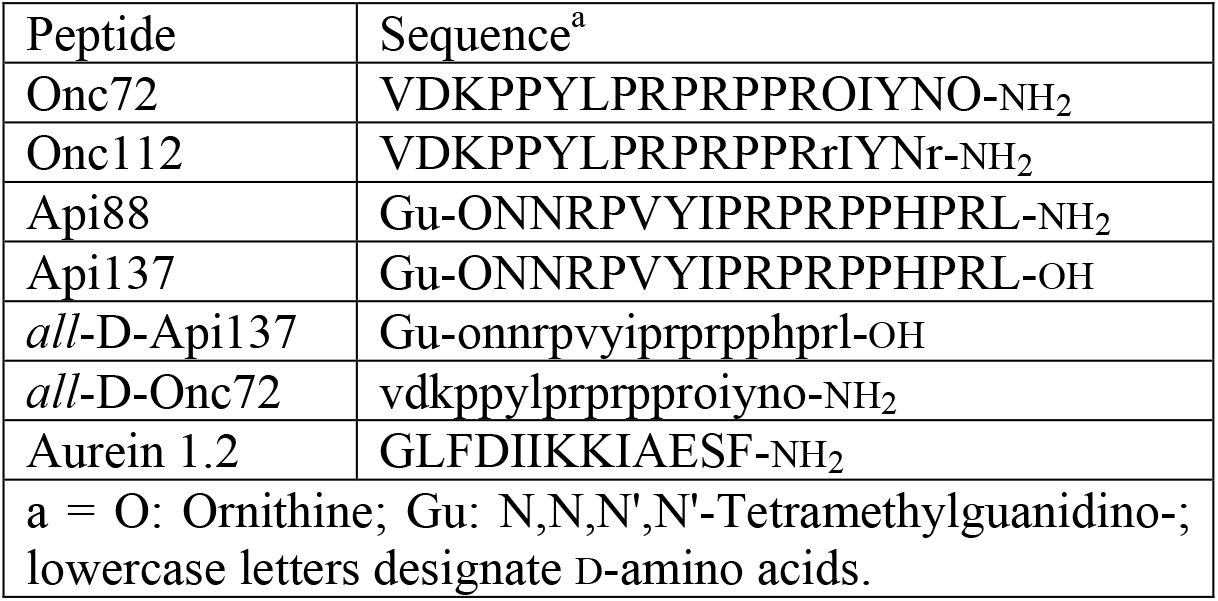
Designation and sequence of antimicrobial peptides.

Here we investigate how the oncocin and apidaecin PrAMPs activity to *P. aeruginosa* and *E. coli* are effected in environments that have varying metal ion concentrations and whether these PrAMPs are able to exert activity that effects the cell envelope of each bacteria.

## Methods and Materials

### Peptide synthesis

Peptides were synthesized by Fmoc/^*t*^Bu chemistry by using either *in situ* activation with *N,N’*-diisopropylcarbodiimide (DIC) in the presence of 1-hydroxybenzotriazole (HOBt) or 2-(1H-benzotriazol-1-yl)-1,1,3,3-tetramethyluronium hexafluorophosphate (HBTU) with *N,N*- diisopropylethylamine (DIPEA)^4, 12^. The solid supports used were Rink amide 4-methylbenzhydrylamine (MBHA) and 4-benzyloxybenzyl alcohol resins (Wang MultiSynTech) to obtain C-terminal amides and free acids, respectively.

The *N,N,N’,N’*-tetramethylguanidino group was incorporated onto the unprotected N-terminus by using 10 equivalents of HBTU and DIPEA or *N*-methylmorpholine in dimethylformamide^4, 12^. The final peptide resins were washed with dichloromethane, dried, and cleaved with trifluoroacetic acid (TFA) containing 12.5% (vol/vol) of a scavenger mixture (ethanedithiol, *m*-cresol, thioanisole, and water; 1:2:2:2 [vol/vol/vol/vol]). After 2 h, the peptides were precipitated with cold diethyl ether, washed twice with cold diethyl ether, dried, and purified with a C_18_-phase column by using a linear aqueous acetonitrile gradient in the presence of 0.1% (vol/vol) TFA (Jupiter C18 column; 21.2-mm x 250-mm length, Phenomenex Inc., Torrance, CA). The peptide purities were analysed by reverse-phase high-performance liquid chromatography and masses were confirmed by matrix-assisted laser desorption/ionization–time-of-flight mass spectrometry (MALDI-TOF MS; 4700 Proteomic Analyzer; AB Sciex, Darmstadt, Germany). Peptide designation, sequence and observed masses are described in table 1.

### Bacterial growth

*Pseudomonas aeruginosa* (ATCC 47085) and *Escherichia coli* strains (ATCC 25922) from the Melbourne Dental School culture collection were cultured from freeze dried stocks and maintained on horse blood agar plates supplemented with 10 % (v/v) lysed horse blood. *P. aeruginosa* or *E. coli* bacterial colonies were used to inoculate Luria Bertani (LB) and grown aerobically at 37°C. Batch culture growth was monitored at 650 nm using a spectrophotometer (model 295E, Perkin Elmer) and culture purity was routinely checked by Gram stain and by colony morphology. For antimicrobial assays *P. aeruginosa* and *E. coli* were grown planktonically until late exponential phase of growth and harvested by centrifugation (8,000×*g*, 30 mins). Viability and bacterial cell number/mL were determined by flow cytometry using a Cell Lab Quanta SC flow cytometer (Beckman Coulter Pty Ltd, NSW, Australia) and a LIVE/DEAD BacLight™ Bacterial Viability Kit, according to the manufactures’ instructions (Life Technologies Pty Ltd, NSW, Australia).

### Antimicrobial and membrane potential assays

Bacteria (*P. aeruginosa* or *E. coli*; 2.5 × 10^5^/100 μL) in LB media were incubated aerobically, 37 °C, with varying concentrations (512, 256, 128, 64, 32, 16, 8, 4, 2, 1, 0.5, 0.0 μg/mL) of AMP (Api88, Api137, Onc72 or Onc112) for 90 min, final assay volume 200 μL (equating to 1.25 × 10^6^ cells/mL). Following incubation, Minimal Inhibitory (MIC) and Minimal Bactericidal Concentrations (MBC) were determined by turbidometric assay and Colony Forming Unit (CFU), as previously described^23^ using LB media or 33% v/v Tryptic Soy Broth (TSB). For determination of the Minimal-membrane Disrupting Concentration (MDC) Syto9 (3.34 mmol/L) and Propidium Iodide (PI, 50 μg/mL) were added to the bacterial samples and analysed by flow cytometry using a Cell Lab Quanta SC flow cytometer (Beckman Coulter Pty Ltd, NSW, Australia) for the proportion of live (Syto9 +, FL1) and dead bacteria (PI +, FL3) using standard flow cytometry settings and MDC calculated as previously described^23^.

Membrane potential was determined using the BacLight™ Bacterial Membrane Potential Kit (Life Technologies, NSW, Australia) according to the manufacturer’s instructions. Briefly, following incubation with AMPs, 10 µL of 3 mmol/L 3,3′-diethyloxacarbocyanine iodide (DiOC_2_(3)) was added to the bacterial samples and incubated for 30 mins at 37°C. Membrane potential was determined by flow cytometry as the proportion of red-DiOC_2_(3) (FL3) to green-DiOC_2_(3) (FL1) mean fluorescence intensity (MFI) of bacterial cells; FL3-MFI/FL1-MFI using a Cell Lab Quanta SC flow cytometer (Beckman Coulter Pty Ltd, NSW, Australia). Bacteria incubated with media alone or in the presence of carbonyl cyanide 3-chlorophenylhydrazone (CCCP; 10 µL of a 0.5 mmol/L stock solution, 30 mins, RT) were used to determine the membrane potential of untreated bacteria and a depolarised membrane, respectively.

Antimicrobial activity and membrane potential are reported as the average ± standard deviation of three biological replicates assays where all assay concentrations had two technical replicates.

### Outer membrane permeabilisation assay, (N-phenyl-1-napthylamine (NPN) uptake assay)

The ability of PrAMPs to permeabilise the outer membrane of *P. aeruginosa* was determined by using the NPN uptake assay as previously described^24^. Briefly, *P. aeruginosa* (ATCC 47085) cells were washed and resuspended in HEPES buffer (5 mmol/L HEPES, 5 mmol/L glucose, pH 7.4) which contained 170 mmol/L NaCl, 30 mmol/L NaCl, and either 1.0 mmol/L MgCl_2_ or 5 mmol/L CaCl_2_. Pr-AMPs were formulated with the respective HEPES buffer and serially diluted in 96 well plate. NPN was directly added to the bacteria suspension to give a final concentration of 10 μmol/L, the bacteria/NPN solution was immediately added to the serially diluted AMP, and the fluorescence recorded (excitation = 355 nm, emission = 405 nm) every 30 secs for 10 mins and then every 10 mins over 1 hour using a Wallac Victor 2V Multilabel Microplate Reader 1420 (Perkin-Elmer, NSW, Australia). Buffer was used as the negative control and polymyxin B (10 μg/mL) was used as a positive control due to its strong outer membrane permeabilising properties. NPN uptake is reported as fluorescence (405 nm) using the equation: F_obs_ – F_0_, where F_obs_ = NPN fluorescence of bacteria + AMP, F_0_ = initial NPN fluorescence of bacteria in the absence of AMP. All values had background fluorescence subtracted: fluorescence of HEPES + NPN minus fluorescence of HEPES without NPN.

## Statistical analysis

The data obtained were determined to be normally or non-normally distributed using homogeneity of variances and assessed using Levene’s test (GraphPad Prism). Statistical analysis was performed using a one-way classification of ANOVA and Student’s t-test (two-tailed), where differences were regarded as statistically significant with probability P < 0.05. The magnitude and direction of differences were determined using effect size (Cohen’s d) where a positive or negative d indicates the direction of the effect and the size of the effect being; large where d>0.8, medium where d<0.8 and >0.5 and small where d<0.5 and >0.2 and no effect where d<0.2.

## Results and Discussion

### Antimicrobial activity of PrAMPs against *P*. *aeruginosa* and *E*. *coli*

*P. aeruginosa* or *E. coli* were incubated at varying concentrations of PrAMPs Api88, Api137, Onc72, and Onc112 and a control membrane lytic antimicrobial peptide (aurein 1.2) and the MIC, MBC and MDC determined (Table 2). All of the AMPs had antimicrobial activity against *E. coli* but had weak or no antimicrobial activity against *P. aeruginosa*. For *E. coli* the MBC’s of Onc72 and Onc112 were significantly (p < 0.05) higher than the MIC, indicating that these AMPs are primarily bacteriostatic. Api88 and Api137 had significantly (p < 0.05) lower MIC’s and MBC’s than the oncocin peptides against *E. coli*, with Api88 having greater activity than Api137, which is consistent with previously reported activity of these peptides^4^. The equivalent MIC and MBC induced by either Api88 or Api137 indicates that the apidaecin peptides are bactericidal against *E. coli*. As can be seen in Figure 1, none of the PrAMPs resulted in an increase in PI+ *E. coli* cells, above the no AMP control, indicating that the peptides do not induce inner membrane disruption. This is in contrast to the membrane lytic peptide aurein 1.2, which induced inner membrane disruption as indicated by the increase in PI+ *E. coli* cells compared to the no AMP control. For aurein 1.2 MDC as well as MIC and MBC could be determined which were all equivalent indicating that aurein 1.2 is bactericidal and lytic against *E. coli* (Table 2).

**Table 2.** Antimicrobial activity of PrAMPs, aurein 1.2, and Ampicillin against *E. coli* and *P. aeruginosa* in LB medium.

|  | Antimicrobial peptide activity (µg/mL ± SD)<sup>a</sup> |  |  |  |  |  |
| --- | --- | --- | --- | --- | --- | --- |
|  | *E. coli* |  |  | *P. aeruginosa* |  |  |
|  | MBC<sup>b</sup> | MIC<sup>b</sup> | MDC<sup>b</sup> | MBC<sup>b</sup> | MIC<sup>b</sup> | MDC<sup>b</sup> |
| Onc72 | 140.1 ± 17.2 | 57.7 ± 19.8 | > 256 | > 256 | > 256 | > 256 |
| Onc112 | 39.5 ± 2.4 | 18.9 ± 1.8 | > 256 | 80.9 ± 16.4 | 81.5 ± 13.6 | > 256 |
| Api88 | 5.3 ± 1.9 | 9.1 ± 1.4 | > 256 | > 256 | > 256 | > 256 |
| Api137 | 9.2 ± 1.2 | 13.9 ± 3.3 | > 256 | > 256 | > 256 | > 256 |
| Aurein 1.2 | 26.1±1.7 | 24.9±2.2 | 25.7±3.4 | ND<sup>c</sup> | ND<sup>c</sup> | ND<sup>c</sup> |
| Ampicillin | ND<sup>c</sup> | 4.83 | ND<sup>c</sup> | ND<sup>c</sup> | > 32 | ND<sup>c</sup> |

a = antimicrobial activity is reported as the average ± standard deviation of two biological replicates with two technical replicates.
b = MBC = Minimal Bactericidal Concentration; MIC = Minimal Inhibitory Concentration; MDC = Minimal-membrane Disrupting Concentration.
c = Not determined.

**Figure 1.**
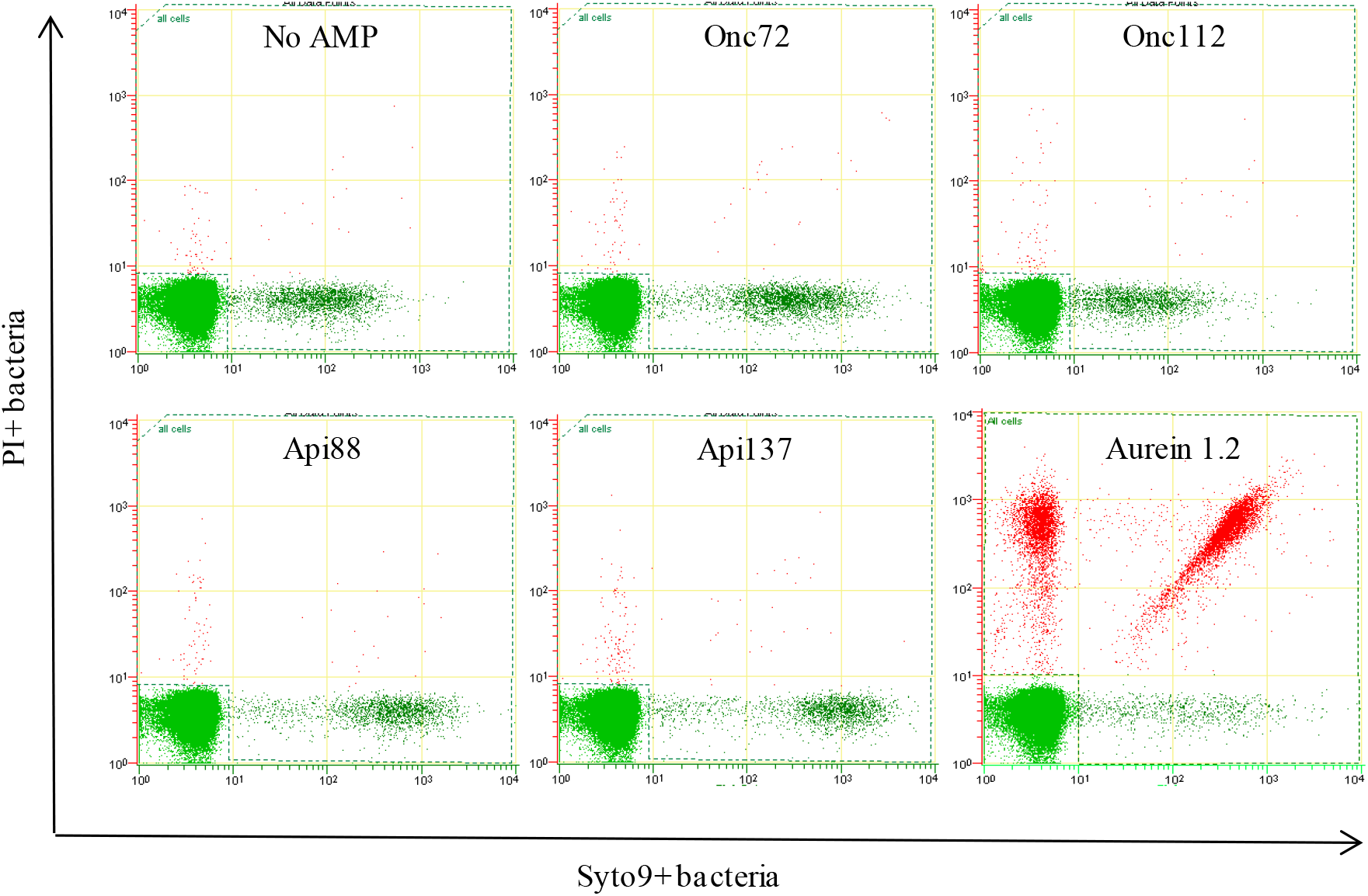
Flow cytometry dot plots showing the relative membrane disruption of *E. coli* incubated with antimicrobial peptides at their respective MIC concentrations.

Although the oncocin and apidaecin peptides were effective against *E. coli*, their activity is higher in previous reports and have also been shown to be effective against *P. aeruginosa* which contrasts with our data^9, 12, 13, 25^. A point of difference between our current work and previous antimicrobial assays is the use of ion-rich LB media in this study compared to diluted Müller Hinton Broth (MHB, 25%) or TSB (33%). To investigate if activity could be improved towards *E. coli* and *P. aeruginosa* we conducted the same assays in 33% v/v TSB media, in line with previous reports^9, 12, 13, 25^. There was a significant improvement in activity towards *E. coli* for both apidaecin and oncocin derivatives (Table 3). Api88 and Api137 still had lower MICs and MBCs than Onc72 or Onc112 in the 33% TSB media indicating they are more active against *E. coli*. Intriguingly, the mode of action for both oncocin peptides against *E. coli* is different between the two media, changing from being bacteriostatic in LB media to bactericidal in 33% TSB. In the 33% TSB media, both the oncocin and apidaecin peptides were able to kill *P. aeruginosa* (Table 3). Unlike the activity for *E. coli* the oncocin peptides had much lower MICs and MBCs against *P. aeruginosa* than the apidaecin peptides, with Onc112 being the most active. For both the oncocin and apidaecin peptides the MICs and MBCs were equivalent, indicating that they are all bactericidal against *P. aeruginosa*. It is known that a mechanism by which *P. aeruginosa* protects itself from antimicrobials is by stabilising the outer-membrane and lipopolysaccharide (LPS) through crosslinking with free cations, such as Ca^2+^, Mg^2+^ or Na^+26-28^. Another mechanism by which *P. aeruginosa* reduces antibiotic/AMP penetration is that the primary porin on its surface OprF is predominately in the closed position, restricting outer membrane permeability by 12-100-fold lower than in *E. coli*^29, 30^. The improved activity of both oncocin and apidaecin peptides against *P. aeruginosa* in the lower ion media indicates that AMPs can be highly effective at killing *P. aeruginosa* if they are able to penetrate the outer membrane. To investigate whether the apidaecin and oncocin peptides penetrate the outer membrane of *P. aeruginosa* via an LPS/lipid route more than a porin route we conducted an outer membrane penetration assay in low and high ion salt concentrations. A major difference between LB media and 33% TSB media (or cationic MHB) is the availability of Na^+^, Ca^2+^ and Mg^2+^ ions. To investigate if these metal ions prevented PrAMPs to penetrate the outer membrane we used the NPN uptake assay, whereby small disturbances in the outer membrane can be observed by the insertion of NPN in the outer membrane resulting in fluorescence. Assays were conducted in HEPES buffer with the relevant amounts of Na^+^, Ca^2+^ and Mg^2+^ ions as chloride salts, equivalent molarity in each media. In conditions that mimicked 33% TSB media whereby the concentrations of Na^+^, Ca^2+^ and Mg^2+^ ions are 30 mmol/L, 0 mmol/L and 0 mmol/L, respectively, all of the active PrAMPs were able to permeabilise the outer membrane of *P. aeruginosa* (Figure 2A, 2B, 2C). Interestingly, in the low salt ion buffers the all-D-Onc72 was able, albeit weakly, to permeabilise the outer membrane (Figures 2A, 2B, 2C). In conditions that mimicked LB media, whereby the concentrations of Na^+^, Ca^2+^ and Mg^2+^ ions were 170 mmol/L, 5 mmol/L and 1.0 mmol/L, respectively, (these concentrations also mimic the levels in mammalian tissue and body fluids) all of the active PrAMPs had no capacity to permeabilise the outer membrane, with the exception of Onc72 and Onc112 in the presence of 1.0 mmol/L MgCl_2_ which had a slight ability to permeabilise the membrane. The control, Polymyxin was unaffected by 170 mM NaCl, had reduced capacity in the presence of MgCl_2_ and very slight activity in the presence of CaCl_2_ (Figure 2D).

**Table 3.**
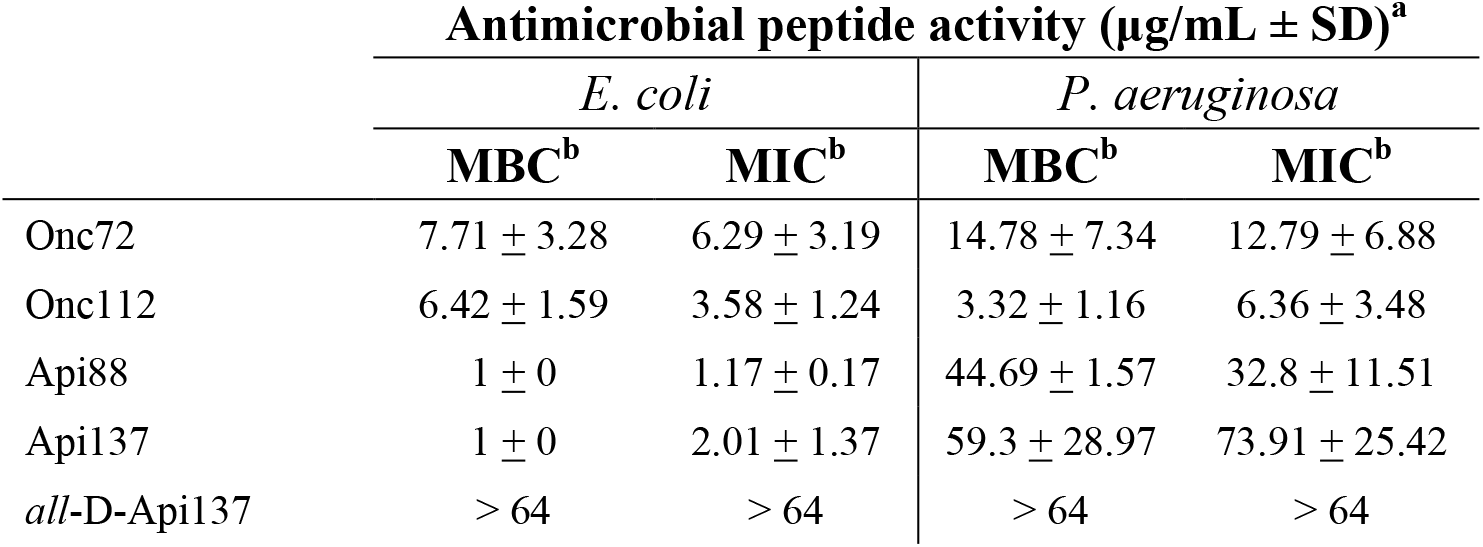
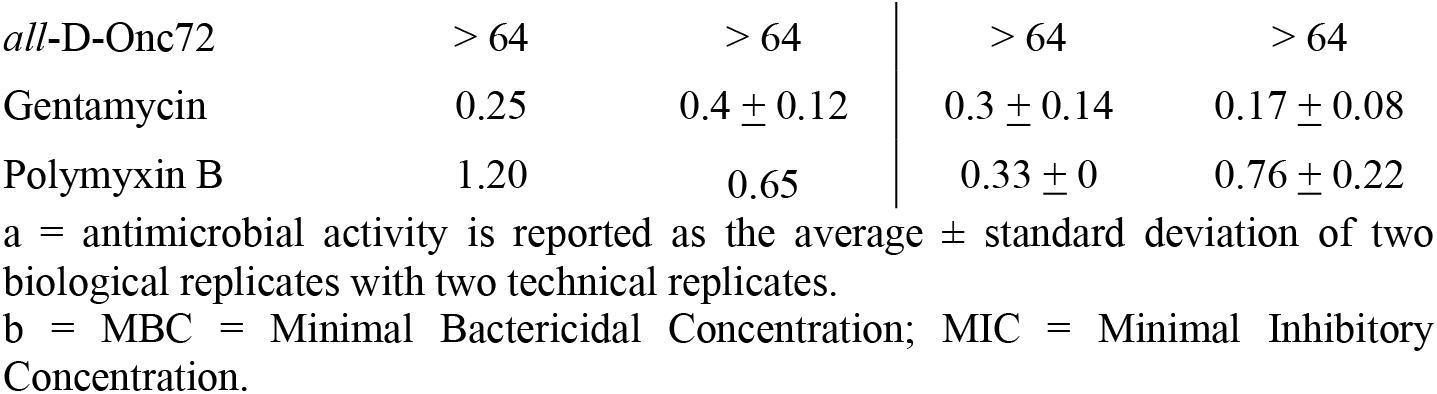
Antimicrobial activity of PrAMPs and antibiotics against *E. coli* and *P. aeruginosa* in 33% TSB media.

**Figure 2.**
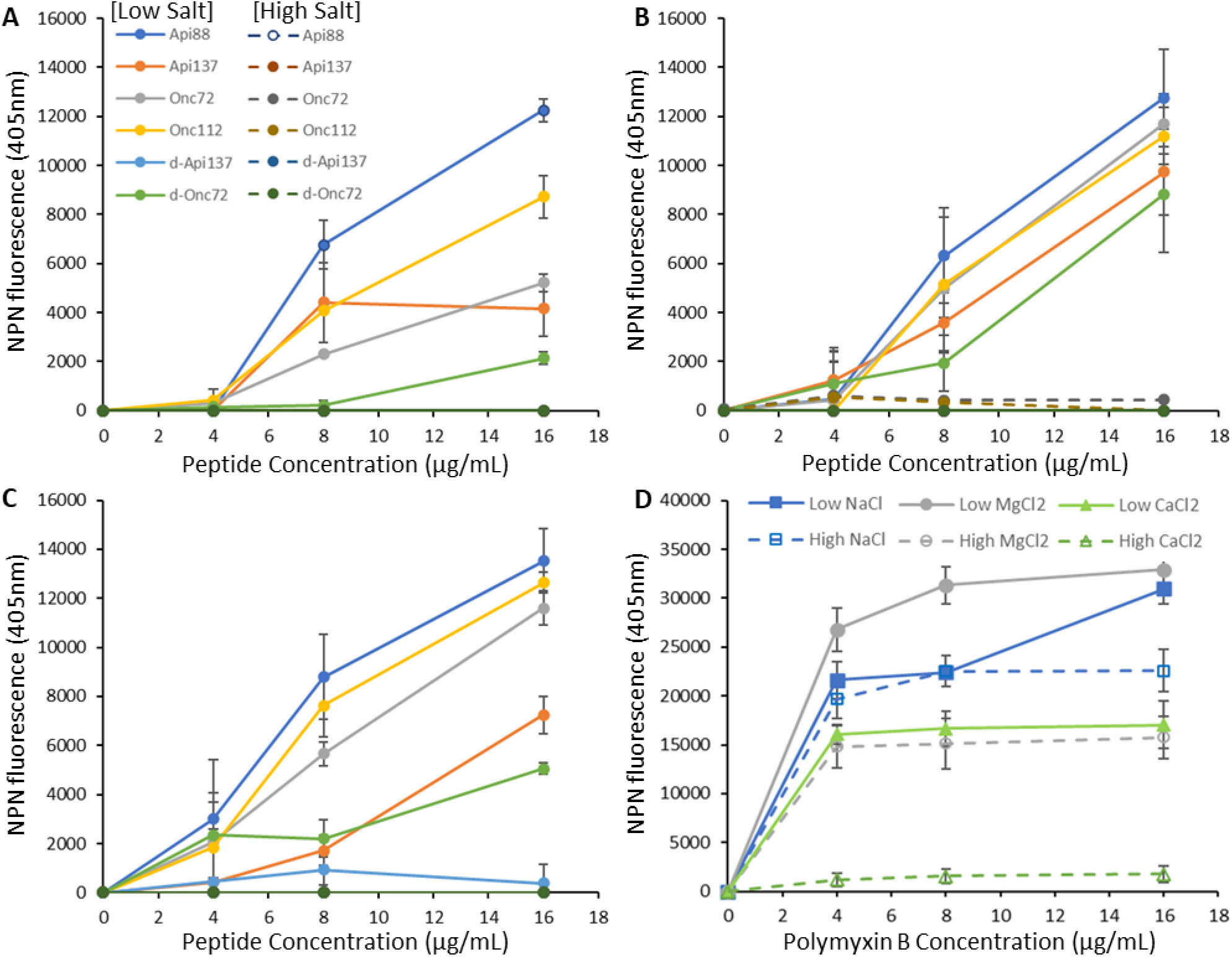
Outer membrane permeation of *P. aeruginosa* by PrAMPs in low salt and high salt buffers. (A) Permeation of *P. aeruginosa* outer membrane in low (30 mmol/L) and high (170 mmol/L) NaCl HEPES buffer by PrAMPs, (B) Permeation of *P. aeruginosa* outer membrane in low (0 mmol/L) and high (1.0 mmol/L) MgCl_2_ HEPES buffer by PrAMPs, (C) Permeation of *P. aeruginosa* outer membrane in low (0 mmol/L) and high (5 mmol/L) CaCl_2_ HEPES buffer by PrAMPs and (D) Permeation of *P. aeruginosa* outer membrane in low and high salt-adjusted HEPES buffer by Polymyxin B. Data expressed as the mean and standard deviation of NPN fluorescence at 405 nm of three biological replicates of two technical replicate assays.

### PrAMPs alter the membrane potential, size and optical density of *E*. *coli* cells

Api88, Api137, Onc72, and Onc112 have been shown above to kill or inhibit growth of *E. coli* without inducing membrane lysis. However, each of the PrAMPs did have a dramatic effect on the size and optical density of the *E. coli* cell population, as can be observed in the side-scatter v electoral volume flow dot blots (Figure 3). All peptides decreased the *E. coli* cell optical density, although not as strong as compared to aurein 1.2 (Figure 4A). Interestingly, the peptide concentration at which a decrease in the *E. coli* cell population optical density is significantly (p < 0.05) different from untreated *E. coli* cells correspond to the MIC for each of the PrAMPs (Figure 4A). Api88 and Api137 resulted in an increase in the size/cell volume of *E. coli* cells in a similar manner to aurein 1.2, up to a maximal volume increase from untreated cells of +20% (Figure 4B). At low peptide concentrations, Onc72 increased the volume of *E. coli* cells but above 32 μg/mL the cell volume returned to normal. Onc112 did not significantly alter *E. coli* cell volume at any concentration tested. The alteration of the optical density and size of *E. coli* cells suggests that although the PrAMPs do not disrupt the cytoplasmic membrane as determined using propidium iodide they do cause a flow of water into the cell. This could be the result of membrane thinning or pore formation that is smaller than the radius of propidium iodide (thus indicating no pore formation), this phenomenon of membrane permeabilization was found for another proline-rich peptide chex-Arg20^31^.

**Figure 3.**
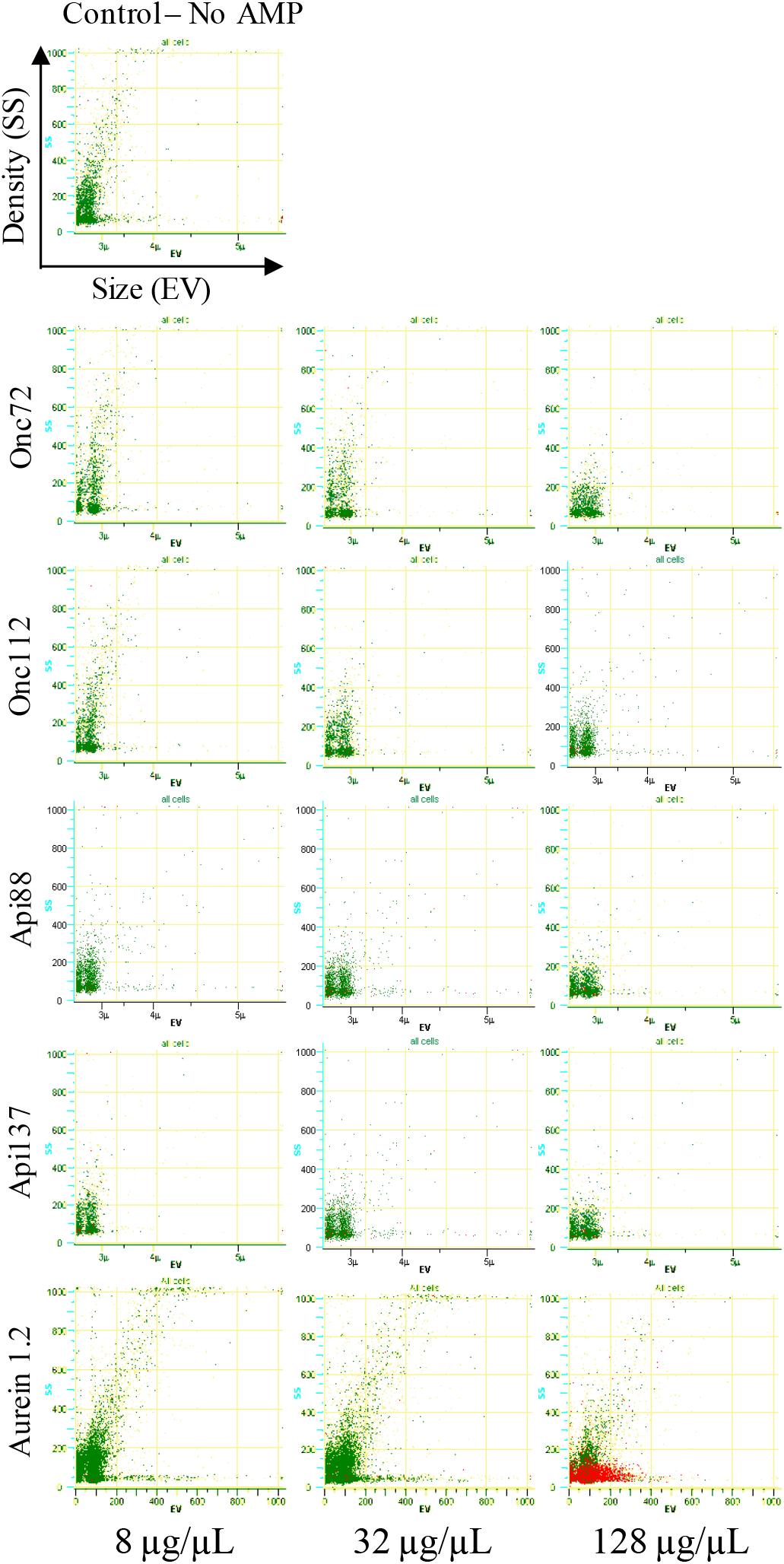
Size and optical density of *E. coli* incubated with antimicrobial peptides at increasing concentrations (Representative flow cytometry dot plots).

**Figure 4.**
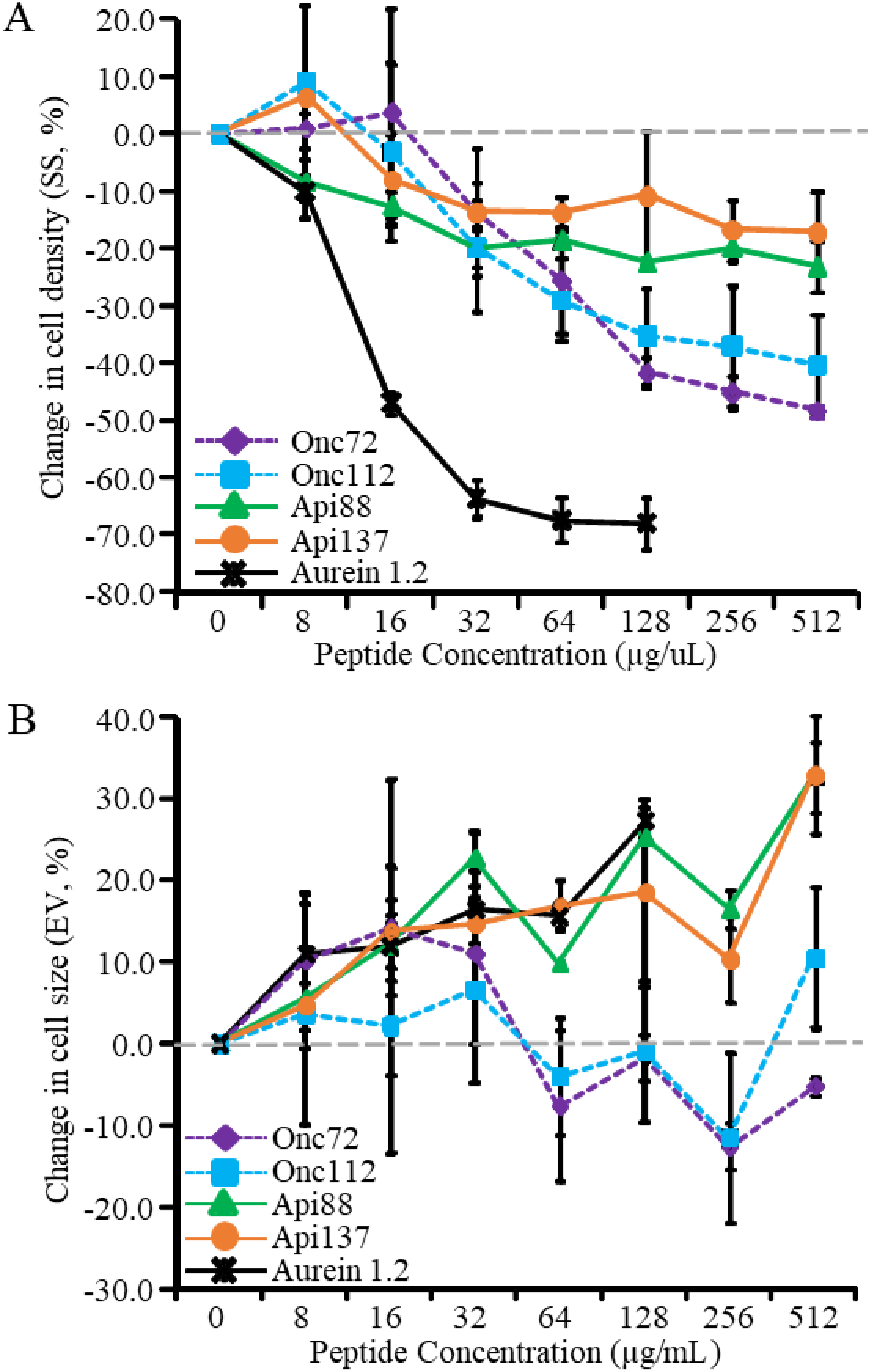
Percent change in *E. coli* cell size and optical density with peptide concentration (A) *E. coli* cell population optical density and (B) *E. coli* cell population size. Dotted grey line represents zero percent change.

To investigate whether the inner membrane of *E. coli* is altered by the PrAMPs, we explored the change in membrane potential when *E. coli* was incubated with each active PrAMP at varying MIC equivalent concentrations (0.5 x, 1.0 x and 2.0 x the MIC of each peptide). Figure 5 shows representative flow cytometry dot plots of DiOC_2_(3) positive *E. coli* cells treated with PrAMPs. Initially *E. coli* cells were incubated with and without the proton ionophore CCCP, which resulted in distinct DiOC_2_(3)+ *E. coli* cell populations; a red fluorescent DiOC_2_(3)+ *E. coli* cell population in the absence of CCCP and a green fluorescent DiOC_2_(3)+ *E. coli* cell population in the presence of CCCP (Figure 5). These distinct DiOC_2_(3)+ *E. coli* cell populations were used to provide flow cytometry gates to distinguish proton gradient intact/altered (polarised, Gate A) and eliminated (depolarised, Gate B) membrane potential populations. All of the PrAMPs resulted in a small but significant increase in the green fluorescent DiOC_2_(3)+ *E. coli* cell population. However this increase was not concentration dependent, as no significant difference was observed in the green fluorescent DiOC_2_(3)+ *E. coli* cell population with increasing peptide concentration. In contrast, the membrane lytic peptide aurein 1.2 resulted in a significant increase in the green fluorescent DiOC_2_(3)+ *E. coli* cell population which increased with peptide concentration. This indicates that the PrAMPs induce a limited depolarisation of the bacterial membrane, whereas, aurein 1.2 induced considerable concentration-dependent depolarisation of the membrane of *E. coli*. Interestingly all of the peptides resulted in a shift in the red fluorescent DiOC_2_(3)+ *E. coli* cell population compared to untreated *E. coli* cells, indicating that membrane polarity is altered by the PrAMPs. By analysing the proportion of red and green fluorescent DiOC_2_(3)+ *E. coli* cell populations it can be clearly seen that all of the PrAMPs induce significant (p < 0.05) hyperpolarisation in membrane potential of *E. coli* cells at each peptide concentration tested (Figure 6). This suggests that the membrane becomes more negatively charged in the in the presence of the PrAMPs. In contrast, aurein 1.2 resulted in significant (p < 0.05) depolarisation of *E. coli* cell membrane (Figure 5), i.e., the membrane becomes less negatively charged. The extent of hyperpolarisation induced by Api88 and Api137 decreased with increasing peptide concentration, whereas the level of hyperpolarisation induced by Onc72 and Onc112 was not affected by peptide concentration. The primary mode of action for many PrAMPs, including the apidaecin- and oncocin-derived peptides, is the prevention of protein translation by binding to the intracellular target the 70S ribosome, but PrAMPs are also known to bind other internal targets such as DnaK, indicating secondary modes of action^15, 32^. In a study using another PrAMP that strongly binds to 70S ribosome, ARV-1502 (Chex1-Arg20)^33^, it was found that it localised to the membrane of *E. coli* and induced a hyperpolarised state, without inducing membrane disruption^34^. Hyperpolarised states are a result of cation efflux or anion influx across a membrane and in a subsequent study, it was shown that Chex1-Arg20 was able to induce anion influx in a LUV lucigenin assay^31^. The similarities in the secondary mode of action of membrane hyperpolarisation between Chex1-Arg20 and the apidaecin-and oncocin-derived PrAMPs, without inner membrane disruption, suggests that Api88, Api137, Onc72 and Onc112 may also act on the inner membrane resulting in anion influx. The reduction in the cell optical density and an increase in volume (cell size) of *E. coli* as shown in figures 4 and 5 are suggestive of an influx of external fluid caused by the studied PrAMPs, and that they result in a weakened membrane. Collectively our data and previous reports indicate that PrAMPs have a multimodal mechanism of action, a primary one of binding to 70S ribosome and preventing protein synthesis and then a set of secondary ones of binding to DnaK and inner membrane interaction resulting in hyperpolarisation a mechanism known to result in cell death.

**Figure 5.**
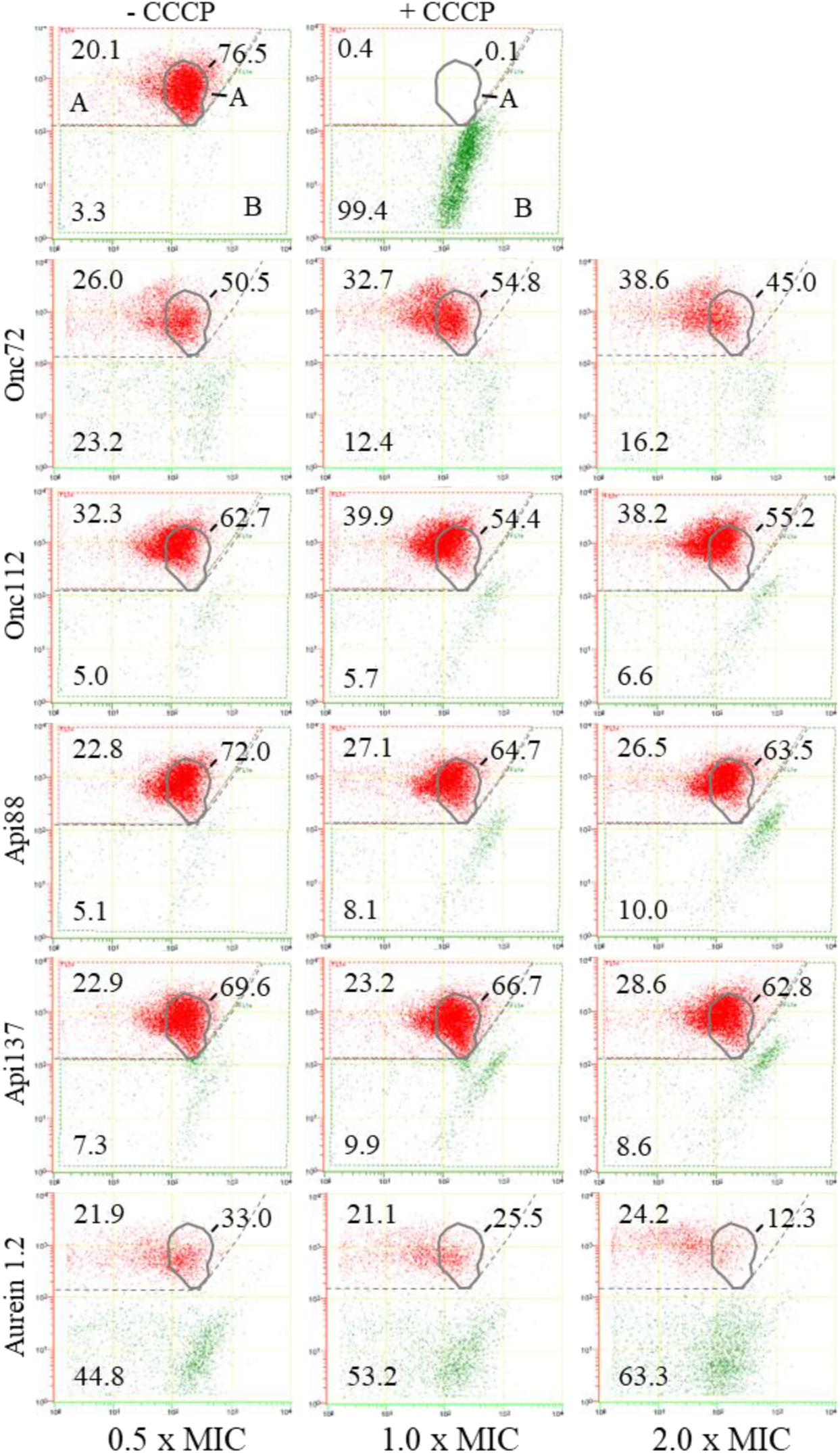
Membrane potential of *E. coli* incubated with antimicrobial peptides at their respective MIC concentrations. A. DiOC_2_(3) red fluorescence indicating an intact/active proton gradient, B. DiOC_2_(3) green fluorescence indicating an eliminated proton gradient (depolarised membrane), C. Intact proton gradient of untreated *E. coli* cells. (Representative flow cytometry dot plots).

**Figure 6.**
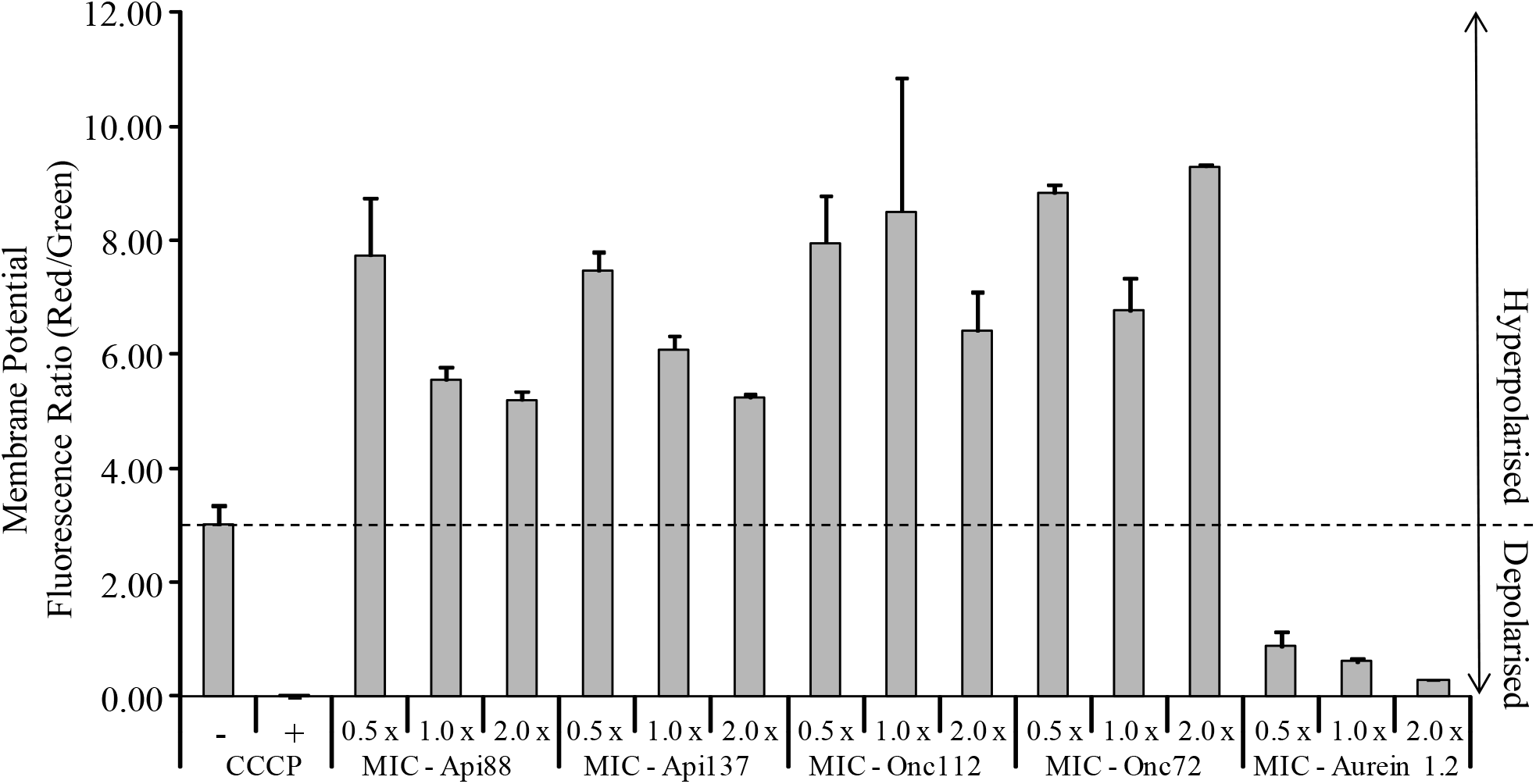
Membrane potential of *E. coli* incubated with antimicrobial peptides at their respective MIC concentrations.

## Conclusion

PrAMPs are considered one of antimicrobial peptides that have great therapeutic potential as they are highly antimicrobial yet have very low toxicity to mammalian cells, are efficacious *in vivo* and can be designed to have improved pharmacokinetic and pharmacodynamic properties. As such it becomes vital to know how PrAMPs act and what conditions effect, either improving or reducing their antimicrobial activity. Here we clearly show that PrAMPs can be highly effective against *E. coli* and *P. aeruginosa* in an environment whereby the bacteria are unable to fortify their outer membrane with Na^+^, Ca^2+^ or Mg^2+^ ions. However, in an environment where Na^+^, Ca^2+^ or Mg^2+^ ions are available such as in body/tissue fluids *P. aeruginosa* is able to fortify its outer membrane PrAMP’s are unable to penetrate through to the inner membrane and cytoplasm. PrAMPs are highly effective against *E. coli* in both ion high and low media, although antimicrobial activity is reduced in high ion media indicating that *E. coli* is also able to fortify its outer membrane but not as effectively as *P. aeruginosa*. The antimicrobial resistance priority list from the WHO places *P. aeruginosa* as critical (priority 1) pathogen for developing therapeutics too^35^. A major difficulty in targeting *P. aeruginosa* is that it is resistant to a wide range of materials, however, our data show that *P. aeruginosa* is highly susceptible to PrAMPs in a low metal ion environment where it is unable to fortify its outer membrane. Thus, a future strategy in targeting *P. aeruginosa* could be a combined PrAMP and metal ion chelator. The results with *E. coli* indicate that PrAMP activity is improved in a low metal ion environment, indicating a PrAMP/chelator strategy may have a broad activity range. Importantly, conducting assays using media that represent bodily fluids may aid in identifying potential clinical targets. Stereoisomerism may play a role in targeting *P. aeruginosa* as Onc112 having 2 D-arginine substitutions for L-arginine in Onc72 and has activity against *P. aeruginosa* in high ion and better activity in low ion environments than Onc72. Inclusion of D-amino acids in a native (L-) peptide is known to improve membrane penetration^36^, and a mixed L- and D-amino acid sequence may be a key parameter as the all D-peptides did not have activity. PrAMPs were shown to permeate the outer membrane of *P. aeruginosa* at which point they are highly efficacious and that they do affect the inner membrane of Gram-negative bacteria causing unregulated ion flow and changes in membrane potential. This secondary mechanism of action for PrAMPs could be a potential additive strategy to be exploited to improve targeting and killing of antimicrobial resistant bacteria.

